# Validation of an engineered Zika virus-like particle vaccine candidate in a mosquito-mouse transmission model

**DOI:** 10.1101/2022.08.08.503125

**Authors:** Maria Vittoria Mancini, Rapeepat Tandavanitj, Thomas H. Ant, Shivan M. Murdochy, Daniel D. Gingell, Chayanee Setthapramote, Piyatida Natsrita, Alain Kohl, Steven P. Sinkins, Arvind H. Patel, Giuditta De Lorenzo

**Affiliations:** MRC-University of Glasgow Centre for Virus Research, Glasgow, Scotland, United Kingdom; Biologicals Research Group, Research and Development Institute, Government Pharmaceutical Organization, Bangkok, Thailand; Department of Clinical Pathology, Faculty of Medicine Vajira Hospital, Navamindradhiraj University, Bangkok, Thailand; Department of Microbiology, Faculty of Medicine, Khon Kaen University, Khon Kaen, Thailand

## Abstract

The primary route of Zika virus (ZIKV) transmission is through the bite of an infected *Aedes* mosquito, when it probes the skin of a vertebrate host during a blood meal. Viral particles are injected into the bite site together with mosquito saliva and a complex mixture of other components. Some of them are shown to play a key role in the augmentation of the arbovirus infection in the host, with increased viremia and/or morbidity. This vector-derived contribution to the infection is not usually considered when vaccine candidates are tested in preclinical animal models. In this study, we performed a preclinical validation of a promising ZIKV vaccine candidate in a mosquito-mouse transmission model using both Asian and African ZIKV lineages. Mice were immunized with engineered ZIKV virus-like particles and subsequently infected through the bite of ZIKV-infected *Ae. aegypti* mosquitoes. Despite a mild increase in viremia in mosquito-infected mice compared to those infected through traditional needle injection, the vaccine protected the animals from developing the disease and strongly reduced viremia. In addition, during peak viremia, naïve mosquitoes were allowed to feed on infected vaccinated and non-vaccinated mice. Our analysis of viral titers in mosquitos showed that the vaccine was able to inhibit virus transmission from the host to the vector.

**Author summary:** Zika is a mosquito-borne viral disease, causing acute debilitating symptoms and complications in infected individuals and irreversible neuronal abnormalities in newborn children. The primary vectors of ZIKV are generally considered to be mosquitoes of the genus *Aedes*, in particular *Aedes aegypti*. Despite representing a significant public health burden with a widespread transmission in many regions of the world, Zika remains a neglected disease with no effective antiviral therapies or approved vaccines to control and prevent infections. The efficacy of several promising candidate vaccines is however under investigation, mainly through artificial infections (i.e. needle-mediated injections of the virus) in animal models, while it is known that components of the mosquito bite lead to an enhancement of viral infection and spread. In this study, we have also included mosquitoes as viral vectors, demonstrating that the ability of a promising candidate vaccine to protect animals against ZIKV infections after the bite of an infected mosquito, and to also prevent its further transmission. These findings represent an additional crucial step for the development of an effective prevention tool for clinical use.

**Graphical abstract:** 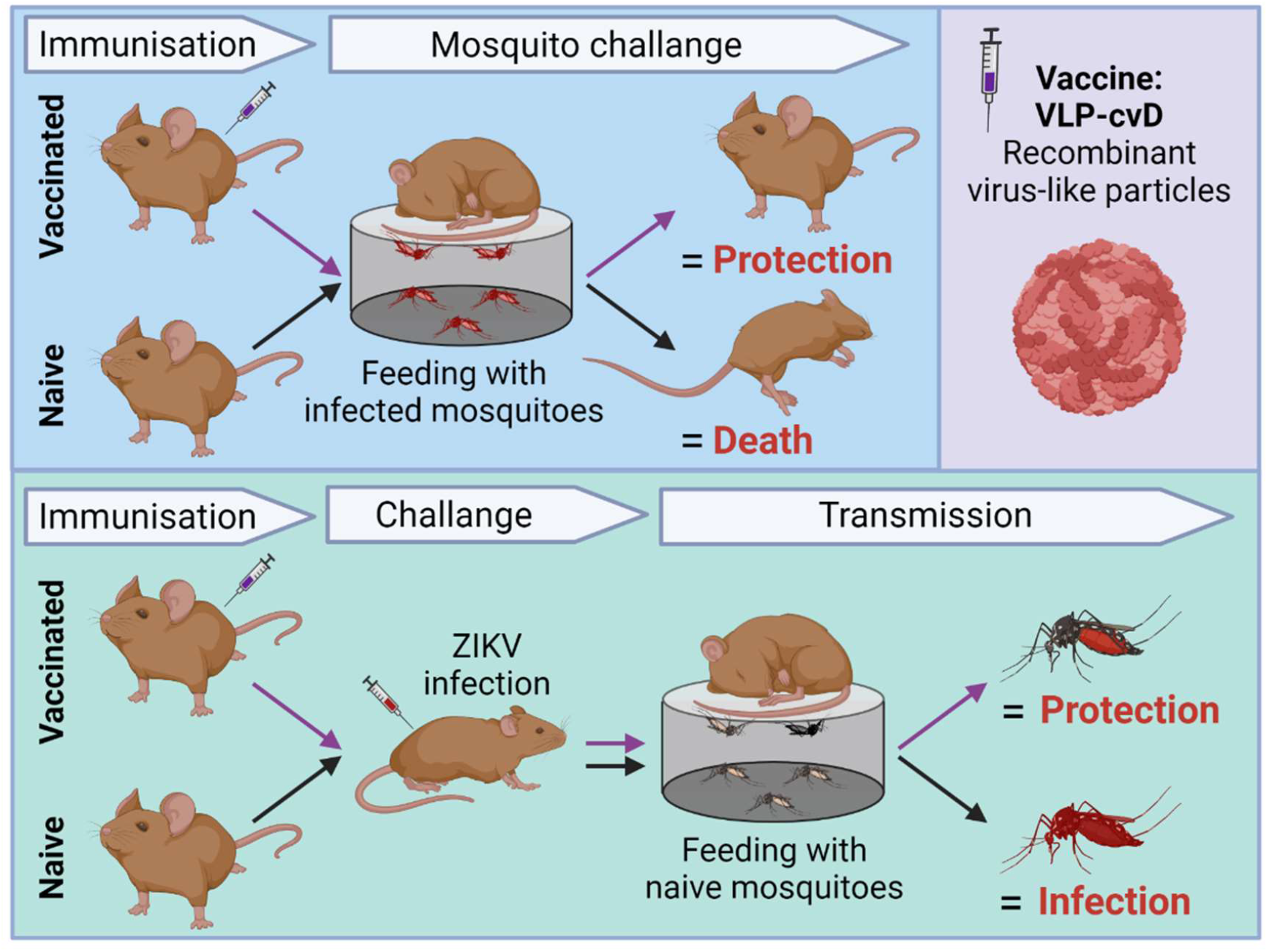

## Introduction

Zika virus (ZIKV) is an arbovirus belonging to the *Flaviviridae* family. Discovered in Africa in 1947, the virus was then introduced in Yap Island (2007), in the South Pacific (2013) and later in Brazil (2015). Historically associated with relatively mild disease, the most recent epidemics linked ZIKV with severe complications, such as neurological disorders (Guillain Barrè syndrome, encephalitis, myelitis)(1, 2) and congenital ZIKV syndrome (CZS) (3, 4). ZIKV is mainly a vector-borne virus but there is strong evidence for sexual (5) and vertical transmission (6). The dramatic spread of the virus in South America during 2016, and the consequent increase in CZS, led the World Health Organisation to declare ZIKV a Public Health Emergency of International Concern. Phylogenetically ZIKV strains are divided in two clades, the ancestral African lineage, and the subsequent Asian lineage which is responsible for the 2007, 2013 and 2015 epidemics. Genetic differences between the two lineages may explain the associated epidemiology. Currently, there is no effective therapy or vaccine available to control ZIKV infection.

During the past few years, several candidate vaccines have been developed and tested in preclinical stages (7-9) and the most promising ones have now moved to clinical trials (10, 11). Preclinical tests for ZIKV vaccine candidates are performed on murine or non-human primate (NHP) models. The standard procedure involves animal immunisation followed by a viral challenge through needle inoculation of laboratory cultivated ZIKV, normally at a dose range of 10^2^-10^4^ plaque forming units (pfu)(12). The most common routes of infection utilised are subcutaneous or intraperitoneal injection, chosen to mimic vector transmission. While such artificial *in vivo* viral challenges using needle injections ensure accurate dose delivery through defined routes, they do not fully replicate the dynamics of vector-mediated transmission of arboviruses, such as ZIKV, into the human or animal host. Human transmission of ZIKV occurs mainly when an infected *Aedes* female mosquito (primarily *Ae. aegypti* and *Ae. albopictus*) feeds on blood to obtain essential nutrients for egg development. During a bloodmeal, a mosquito injects saliva containing ZIKV particles which can infect cells at the bite site and disseminate throughout the vertebrate host. Through the same process, a naïve mosquito can also acquire virus present in the blood stream by feeding on an infected host. To achieve transmission by a mosquito vector, virus present in a bloodmeal must first cross the midgut epithelium and establish an infection in the salivary glands (SG).

Laboratory-cultured virus stocks may include cell components not commonly present in natural infections, and they therefore represent a possible source of artefacts that may impact on the study outcome. In addition, arboviruses naturally replicate in mosquito cells, potentially incorporating vector-specific modifications into virions (e.g., differentially glycosylated surface proteins (13)). Indeed, it has been demonstrated that serial passages in mosquito or mammalian cell lines induce different genetic and phenotypic effects on viral populations (14). It should also be noted that a needle does not inject within the same cellular layers as a mosquito while feeding and probing; moreover, mosquitoes inoculate high doses of virus extra-vascularly and low doses intra-vascularly in a live host (15). Most importantly, viral inoculation occurs when mosquitoes deposit saliva as they probe the skin for a blood meal.

Mosquito saliva is an extremely rich molecular cocktail containing a variety of biologically active effectors that facilitate blood feeding by promoting vasodilatation, and through modulation of host haemostasis and immune response. There is ample evidence that a mosquito bite induces significant effects on viral infectivity and pathogenicity compared to needle-mediated inoculations by a mechanism generically defined as “mosquito-bite enhancement” of infection(16). Mice that received a combined inoculation of the related West Nile virus (WNV) and salivary glands extracts (SGE) showed significantly enhanced viremia compared to those that received the components in distal locations (17, 18). In addition, inoculation of the flavivirus dengue virus (DENV) by mosquitoes correlated with higher and longer viremia than inoculation via subcutaneous, intradermal, or intradermal together with SGE needle injection (19). Similarly, intradermal infection of a bunyavirus, Rift Valley fever virus (RVFV), in conjunction with SGE induced a significant increase in viral titres in blood, brain, and liver, and more severe thrombocyto/leukopenia in mice (20). How saliva creates a favourable environment for the first stages of flavivirus infection remains largely unknown, as only a small number of saliva proteins has been extensively characterized in the host. It is plausible that the mosquito vector contributes to the outcome of infection and its severity by affecting primary target cells, virus dissemination pathways, and immune activation, and by creating a favorable environment suitable for arbovirus replication. Mosquito saliva proteins are known to alter vascular permeability and to augment ZIKV infectivity in mammalian hosts by inhibiting the lymphotoxin-beta receptor (LTbR)-mediated host immune response (21). SGE components are also involved in changes in cytokine and innate signalling profiles, by decreasing the expression of IFNβ/γ, IL-2, and TLR3/7, while increasing IL-4, IL-10, and IL-12 during DENV and WNV, as well as the alphavirus chikungunya virus (CHIKV) infections. The localization and the site of injections impact the type of immune cells encountered by the viral particles. After mosquito bite, T-cell immune responses switch from a Th1 to a Th2 subtype (22-24). The edema formed after a mosquito bite was shown to increase virus retention at the site of inoculation, delay migration to the draining lymph nodes, and facilitate infection of neutrophils (25). Indeed, saliva effectors were shown to lead to the migration of monocytes, neutrophils, eosinophils, and plasma cells to the site of DENV, CHIKV, and WNV (16, 26-29) infection. Interestingly, a salivary gland protein was also found to significantly increase DENV replication in keratinocytes by inhibiting the secretion of antimicrobial peptides and IFNs (30). In addition to the impact on virulence and viral infectivity, mosquito-mediated infection also alters viral tissue tropism in the host: ZIKV-infected non-human primates displayed systemic infections when challenged through mosquito bites, with viral dissemination in the hemolymphatic tissues, female reproductive tract, liver, and kidneys, whereas after subcutaneous needle injection the virus was detected in the cerebrum and the eyes of some individuals (31).

Viral infection routes are not only relevant when investigating virus dissemination dynamics and tropism in hosts, but also for assessing the efficacy of experimental vaccines and therapies against vector-borne pathogens. Virus inoculation through natural vectors includes exposure to factors able to modulate vaccine-induced protective immunity. A promising vaccine candidate against *Leishmania* parasites failed to protect against infected sandfly challenge, in contrast to the initial promising outcome observed following a needle challenge (32).

We previously reported a candidate vaccine comprising ZIKV virus-like particles (VLPs) that have been engineered to display the viral envelope (E) protein locked into a stable dimeric conformation via a disulphide bridge (VLP-cvD)(7). This vaccine conferred protection against both Asian (PRVABC59) and African (MP1751) lineages of ZIKV in a mouse model through the generation of strongly neutralizing antibodies and dramatically reduced virus dissemination to organs, such as brain and testis. Animal challenges, however, were performed with laboratory cultured ZIKV inoculated through a subcutaneous injection. Here, we further validate our candidate vaccine taking into account the three key elements of vector-borne pathogen transmission: the virus, the mammalian host and the mosquito vector. Applying a “mosquito-host” model, we confirm the efficacy of the vaccine to protect from a vector-transmitted ZIKV disease and, by using the same transmission system model, we demonstrate its capacity to also impair the transmission of the virus.

## Results

### Protection against Asian ZIKV strain following vector-mediated infection

We first aimed to evaluate the efficacy of our ZIKV vaccine against the Asian lineage of the virus in a mosquito transmission mouse model. Two groups each of six A129 K/O mice (*IFNα/βR-/-*) received three doses of 2 µg AddaVax-adjuvated VLP-cvD (Vaccinated) or AddaVax in PBS (Control), respectively, by subcutaneous injection (Fig 1A). Separately, one week prior to the end of the immunization schedule, 2-4 days old female mosquitoes were intrathoracically injected with a ZIKV strain belonging to the Asian lineage (PRVABC59). Intra-thoracic injection was preferred to a standard oral feeding with infected blood to ensure that most mosquitoes acquired the virus.

**Figure 1:**
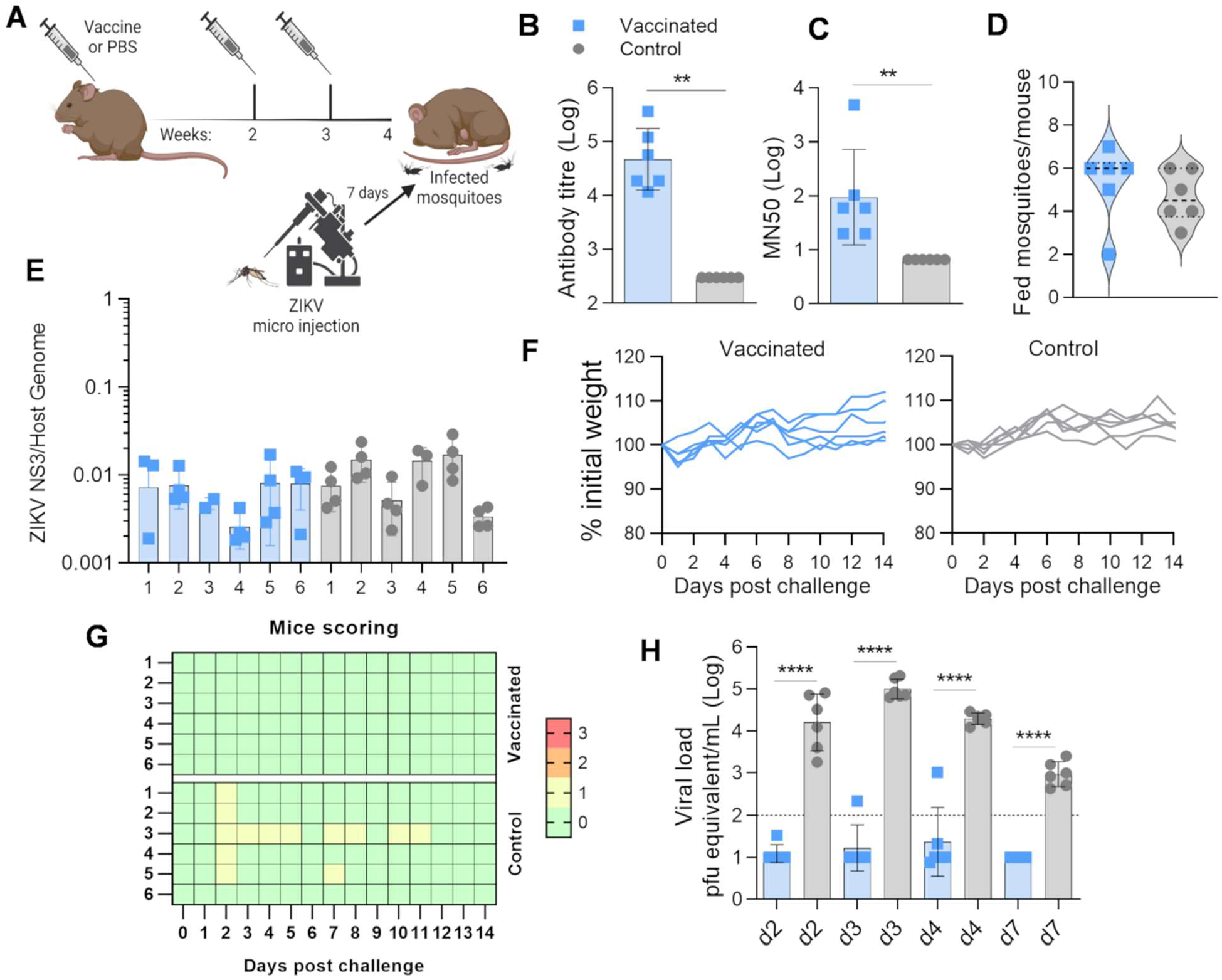
Evaluation of VLP-cvD vaccine efficacy against ZIKV PRVAB59 in a mosquito-mouse transmission model. **A)** Schematic representation of immunisation and challenge procedure, each (n=6) of 4-week-old A129 mice received 3 doses of VLP-cvD or PBS mixed with AddaVax adjuvant. Seven days prior to bite-mediated challenge, mosquitoes were infected with ZIKV by intra-thorax micro injection. **B)** Anti-dimeric E antibody titres of sera collected from animals immunized with VLP-cvD (blue) or PBS (grey). Antibody titres were determined using ELISA plates coated with mono-biotinylated dimeric E. The titre was defined as the maximum dilution that gives a value higher than three-times the value given by the pre-immune sera. Control sera were negative at the lowest dilution (1:900), and their titre was calculated as one-third of that dilution (300). Graph shows three independent experiments represented as Geometric mean with geometric SD. **C)** Neutralization of PRVABC59 ZIKV infection: serially diluted samples of mouse sera were incubated with ZIKV for 1 h before infecting Vero-furin cells. At 72 h post infection, the intracellular levels of E were determined by capture sandwich ELISA, and the percentage infectivity relative to that of the virus alone was calculated. The results were plotted as MN50 values. Graph shows the average of three independent experiments represented as Geometric mean with geometric SD. **D)** Number of fed mosquitoes per mouse at the end of the feeding procedure. Mice were anaesthetized and put on cardboard cups containing 10 infected mosquitoes each. After 20 minutes, mice were removed, mosquitoes anaesthetized by exposure to low temperature, and the engorged mosquitoes counted. **E)** Relative quantification of ZIKV genome copies normalized on the host genome in mosquito carcasses. **F, G)** Animals were weighed (**F**) and scored for clinical signs daily post-challenge (**G**). Legend of scoring system used to monitor animal health following ZIKV challenge: 0 (green) for no signs of distress or disease, 1 (yellow) for one sign of distress, 2 (orange) for two signs of distress or mild disease, 3 (red – humane endpoint) for more than two signs of severe disease or loss of 15% percent of the body weight. **H)** Viral titre in challenged animals. The levels of ZIKV in the serum at days 2, 3, 4, and 7 post infection were quantified by RT-qPCR and the results plotted as equivalent PFU per millilitre. The limit of quantification was estimated to be 100 PFU/ml, indicated by the dotted line. Columns show geometric means from all mice with geometric SD. Assays were performed in triplicate.

Analyses of pre-challenge sera collected (at week 4 post-immunization) from mice confirmed that, unlike the control group, the vaccinated group had high titres of both anti-ZIKV E, and PRVABC59 virus neutralizing antibodies (Figs 1B and C, respectively) (p=0.002, Mann-Whitney). We then performed the virus challenge by allowing the PRVABC59-infected mosquitoes to feed on groups of anesthetized vaccinated and control mice. After feeding, fully engorged mosquitoes were counted to estimate the number of individuals that had a complete meal on each mouse (Fig 1D). An average 40-60% feeding rate was achieved. Total RNA was extracted from the carcasses of the engorged mosquitoes and analysed by RT-qPCR to confirm the presence of ZIKV (Fig 1E). It is not possible to determine the precise dose of virus delivered into the host through mosquito bites, nor is it possible to compare and/or correlate between viral titre in whole mosquito bodies and the effects of the infection on mice. However, a quantitative analysis of ZIKV titres in each engorged mosquito after the blood meal provides an indication of the overall mosquito viral load and showed no difference between the groups (p=0.08, Kruskal Wallis Test).

After the challenge, mice were monitored daily for 14 days for body weight and clinical signs of infection. Surprisingly, we found the overall effect of infection to be generally mild. Excluding the small loss of weight at 1-day post-infection (dpi), which is most likely due to anaesthesia, none of the animals achieved the 10-15% of weight loss typically associated with ZIKV infection (Fig 1F). Likewise, no other clinical signs of infection were observed during the entire monitoring period, with all the vaccinated mice displaying no signs of disease (score 0) and the control mice occasionally showing mild signs of distress (score 1) (Fig 1G). At 2, 3, 4 and 7 dpi, test bleeds were taken, and serum used to quantify the level of ZIKV in the blood by RT-qPCR. Despite the absence of obvious clinical signs, viremia equivalent to up to 10^5^ pfu equivalent/ml was observed indicating an active and consistent infection in all control mice (Fig 1H). In contrast, the viral RNA levels in sera from 3 out of 6 vaccinated mice were below the limit of detection (10^2^ pfu equivalent/ml), whereas a small amount of virus with a peak of only 10^3^ pfu equivalent /ml was present in the serum of the remaining one animal.

Altogether, these findings confirmed the capacity of the VLP-cvD to reduce viremia in mice challenged with mosquito-mediated ZIKV PRVABC59 transmission (p<0.0001, 2way ANOVA). Since no severe symptoms were developed by mice, the assessment of the vaccine-conferred protection against lethal infections was not possible to achieve with this specific ZIKV strain.

The lack of a visible manifestation of symptoms in A129 mice after infection with ZIKV PRVABC59 was unexpected since this strain is commonly used in ZIKV laboratory animal challenge models, usually with reported development of symptoms after an average infection with 10^4^ pfu by subcutaneous injection (7, 33). A similar lack of development of clinical symptoms and lethality was also observed in A129 mice during the initial preliminary infection performed to set up the mosquito-mouse infection model (Sup Fig 1). We reasoned that because AG129 K/O mice (*IFNα/β/γR-/-*) lack interferon I and II receptors, they may be more susceptible to ZIKV infection and pathogenesis. Using groups of six animals each, we compared needle injection and mosquito bite as routes of infection, monitoring them for 14 days as described above. An average of 6 engorged mosquitoes per mouse was measured in the mosquito-challenged group (Sup Fig 2.A), which resulted all positive to ZIKV infection (Sup Fig 2.B). In both cases, ZIKV PRVABC59 was lethal (Sup Fig 2.C), with no apparent differences in clinical scores (Sup Fig 2.D) and weight loss (Sup Fig 2.E) between the groups. Viremia was slightly lower in the needle-administered group at 3 dpi, but the difference was not significant (Sup Fig 2.F). This data confirmed that a vector-mediated PRVABC59 infection supports a lethal infection in the *IFNα/β/γR-/-* mice (AG129) but not in *IFNα/βR-/-* mice (A129), likely due to the enhanced immunodeficiency of the former strain.

### Protection against African ZIKV strain vector-mediated infection

We next tested the efficacy of the VLP-cvD vaccine against the African MP1751 ZIKV strain, reported to induce more severe clinical signs and mortality compared to Asian lineages (34). For direct comparison, we also included in the study an additional group of mice for performing virus challenge via the needle route. Two groups of six A129 mice each were immunized with VLP-cvD and two groups with PBS-control as described above. As expected, the pre-challenge sera from the vaccinated mice (needle and mosquito-challenged) had comparable high levels of anti-E antibodies (p=1, Kruskal Wallis Test) (Fig 2A) and they efficiently neutralized MP1751 ZIKV infection of cultured cells (Fig 2B). Animals in one vaccinated and one control group each were anaesthetized and challenged by infected mosquitoes while the other two groups received a needle inoculation of 100 pfu in 100 µl via the subcutaneous route. Engorged mosquitoes collected from 5 out of 6 animals of each mosquito transmission group were confirmed by RT-qPCR to carry the virus (Figs. 2C and D). As described above, the infected animals were monitored daily for 14 days for weights loss (Fig 2F) and scored for clinical signs (Fig 2G). This time, consistent with the higher virulence of this African lineage ZIKV strain, each individual from the unvaccinated control groups loss weight during the course of infection and exhibited clinical signs that progressively got worst eventually reaching humane endpoint between 6-8 days on average (Fig 2E). In contrast, all vaccinated animals survived mosquito-mediated challenge with no apparent signs of disease. Survival and protection from disease development was excellent also for needle-injected vaccinated mice, except for one individual which lost weight by 15% at day 6 and exhibited clinical signs (Figs. 2F and G). However, upon further analysis the fatality of this animal was considered not to be related to the ZIKV infection and is further discussed below. Interestingly, no engorged mosquitoes were collected from one mouse belonging to the control group infected by mosquitoes; nevertheless, this animal was found to develop the disease, reaching the endpoint 3 days after the rest of the group (from which an average of 3 engorged mosquitoes per mouse were collected) (Fig 2, red arrow). This suggests that unfed/non engorged mosquitoes can also cause a lethal ZIKV infection, most likely a result of virus inoculation during the initial probing phase of blood meal acquisition. At day 3 and 5 post-challenge, mice sera were collected for quantitative analysis of viremia, which showed a significant reduction of virus presence in both vaccinated groups (Fig. 2H) (p<0.0001, 2way ANOVA). The comparison of the viral loads between control groups infected by mosquitoes or by needle displayed a higher, albeit not significant, viremia in mice when the infection was mediated by the invertebrate vector.

**Figure 2:**
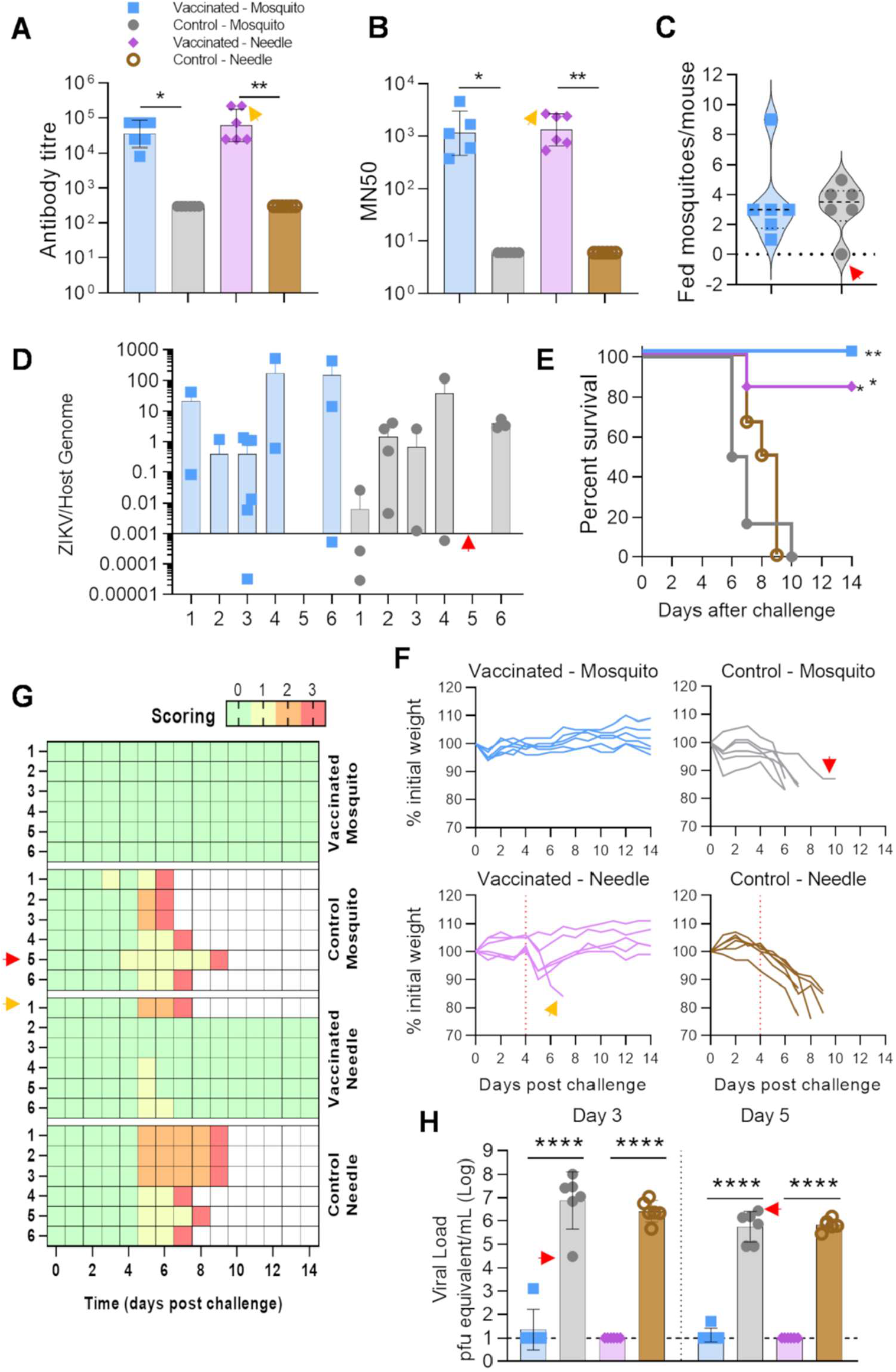
Evaluation of VLP-cvD vaccine efficacy against ZIKV MP1751 in a mouse-mosquito transmission model. **A)** Anti-dimeric E antibody titres of sera collected from animals immunized with VLP-cvD (blue and pink) or PBS (grey and brown). Graph shows three independent experiments represented as Geometric mean with geometric SD. **B)** Neutralization of MP1751 ZIKV infection. The results were plotted as MN50 values, i.e., titres at which 50% neutralization was achieved. Graph shows three independent experiments represented as geometric mean with geometric SD. **C)** Number of fed mosquitoes per mouse at the end of the feeding procedure. **D)** Relative quantification of ZIKV genome copies normalized on the host genome in mosquito carcasses. **E)** Percentage of survival mice in the course of the 14-days challenge, **F)** Mouse scoring for signs of infection. **G)** Animal body weight variations, calculated as percentage of the initial weight. Red line indicated the day of the transmission feeding procedure (day 4). **H)** Viral titre in challenged animals at days 3 and 5. Columns show geometric means from all mice with geometric SD. Quantification performed in triplicates. Dotted line indicates limit of quantification. Red arrow indicates Control – Mosquito mouse 5. Orange arrow indicates Vaccinated – Needle mouse 1.

### Transmission blocking in immunized infected mice

An additional objective of this study was to assess whether our vaccine was able to interfere with ZIKV transmission from an infected mammalian host to the invertebrate vector. We explored this hypothesis using the “vaccinated-needle” and “control-needle” groups of animals described in the previous experiment (Fig. 2F). In compliance with the animal licence regulating this study, where repetitive mosquito feedings are restricted to once a week, the “needle” groups were chosen over the “mosquito”, since the peak of viremia in mice was identified at day 4 post infection using ZIKV MP1751. At the expected peak of viremia vaccinated and control A129 mice were anaesthetized, and naïve mosquitoes were allowed to feed on them (Fig 3A). Blood samples collected at day 3 and 5 after challenge confirmed the high viral load (Fig 3B). As previously described, mice experienced a reduction in body weight on the day after the anaesthesia, the vaccinated group rapidly recovered (Fig 2F – red line) except for one individual, later culled together with the highly infected control group. However, analysis of the serum taken from this individual before and after challenge displayed elevated titre of neutralizing antibodies (Fig 2 1-B, orange arrows) and no viremia, respectively, indicating that most probably the death was not caused by ZIKV, but it was a consequence of anaesthesia (Fig 3B – Vaccinated 1, orange arrow). Engorged mosquitoes were selected and maintained for 14 days to allow the virus to disseminate and reach the salivary glands. An average of 7-9 engorged mosquitoes was collected per mouse (Fig 3C – Day 0), with no significant mortality rate at the end of the incubation period (Fig 3C – Day 14). At day 14, mosquito salivary glands were dissected, and viral load was quantified by RT-qPCR in the carcasses (Fig 3D) and by fluorescent focus assay (FFA) on salivary glands (Fig 3E). Analysis showed that all mosquitoes (43/43, 100%, Fig 3F) fed on infected unvaccinated control mice displayed high disseminated ZIKV load (Fig 3D-E – brown bars). On the contrary, among the mosquitoes fed on infected vaccinated mice, only one (1/32, 3.1%, Fig 3F) (p=<0.0001, Fisher’s exact test) had acquired infection with low virus titre (Fig 3D-E - purple). Thus, vaccination offered an excellent level of protection against viral transmission from the mammalian host to the vector, a critical aspect of arbovirus lifecycle.

**Figure 3:**
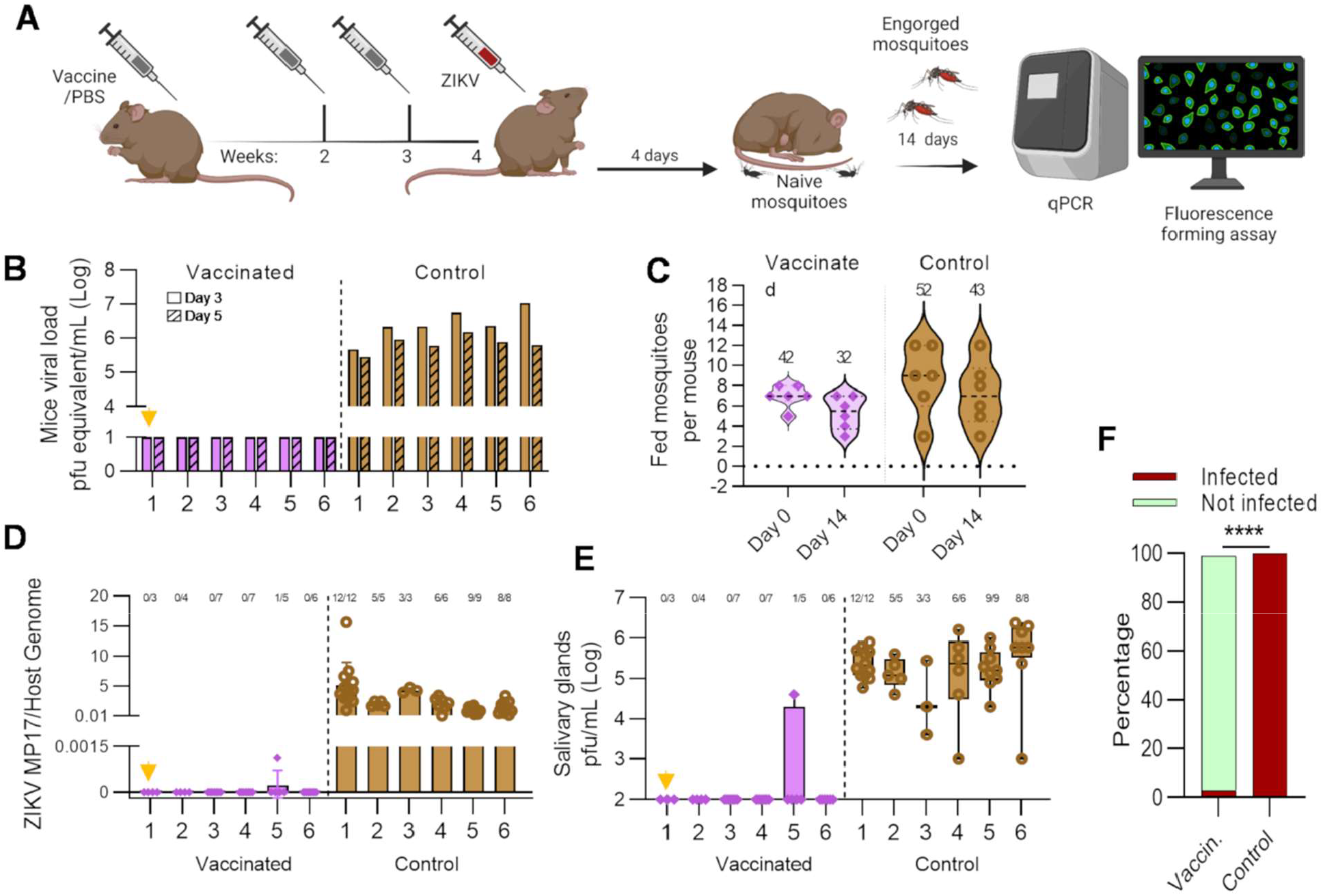
Reverse transmission from mammalian host to invertebrate vector. **A)** Schematic of mouse immunisation, virus challenge and subsequent post-feeding mosquito analysis. Animals correspond to the “needle” groups are already described in Fig 2. **B)** Viral titre in challenged animals at days 3 (solid column) and 5 (stripped column). X-axis indicates ID of each mouse, vaccinated (pink) and control (brown). Quantification performed in triplicates. **C)** Number of mosquitoes that completed a blood meal on VLP-cvD (pink) or PBS (brown) injected mice. Graph shows the number of engorged mosquitoes at the end of the feeding procedure (day 0) and the individual surviving the 2 weeks incubation period (day 14). **D)** Relative quantification of ZIKV genome copies normalized on the host genome from mosquito carcasses. **E)** Quantification of viral titre in homogenized mosquito salivary glands through FFA. Data are represented in Min to max box plot with all points. Orange arrow indicates Vaccinated – mouse 1. **F)** Percentage of infected (red) and not infected (green) mosquitoes 14 days post feeding in vaccinated or control mice. Data are shown as percentage on the total of the analysed individuals.

## Discussion

It is well established that mosquito-derived factors influence arbovirus infection in several ways, with implications on viral titre, tropism, and even disease severity. In this work, we have incorporated a vector-derived virus transmission to a mouse ZIKV model: this represents an epidemiologically relevant mode to study real vaccine efficacy in preventing ZIKV disease and transmission. We confirmed the capacity of our vaccine candidate to protect from the disease in a mosquito-mediated challenge as well as after needle injection. In addition, by developing a mouse-mosquito transmission model, we have also assessed the capacity of the vaccine to inhibit viral transmission from the infected vaccinated mammalian host to the mosquito vector.

Numerous models and techniques may be applied to mimic the enhancement of infection caused by a mosquito bite, spanning from spiking the virus stock with saliva or SGE to the inoculation of virus directly in the site of mosquito bites (spot feeding). These methods are at best artificial proxies and therefore, care needs to be taken when designing transmission blocking assays. Isolation of mosquito saliva is obtained by forcing salivation of females into a glass capillary, often containing mineral oil: the saliva obtained during this artificial collection differs qualitatively and quantitatively from the saliva that is naturally inoculated during a blood meal. SGE, on the other side, is an extract of all the proteins present in the salivary gland tissues, and not just the secreted salivary proteins/effectors that are injected into the host during mosquito probing. We considered these approaches as too prone to experimental artefacts and, thus we opted to replicate the natural mosquito-mediated transmission cycle. This method does not involve collection steps and has the advantage of guaranteeing that the inoculated virus has been propagated directly in the mosquito body, again mimicking the natural transmission setting. However, a caveat is the lack of a quantitative control on the inoculated dose, which may vary according to the volume of saliva inoculated by each mosquito and with the number of attempted probing events. The number of mosquitoes that completed the meal was determined by counting fully engorged females at the end of the pre-set feeding time. While this number considerably varied from mouse to mouse, it did not lead to a comparable variability on the infection progression, with consistent viremia and mortality among the control groups. The number of fed mosquitoes per mouse was not representative of the probing events that occurred. In one instance, no blood-fed mosquitoes were collected from a control mouse (Fig 2, Control mouse 5, red arrow) which, nonetheless, tested positive for infection. This suggests that mere probing can lead to sufficient ZIKV transmission to induce host infection, without the uptake of a full or partial bloodmeal, albeit with delayed viremia and pathogenesis insurgence, compared to controls.

Despite PRVABC59 being a ZIKV strain widely used in *in vivo* and *in vitro* experimental settings, in our hands it did not induce clinical symptoms and lethal infections in A129 mice when inoculated through a mosquito bite, although viremia was detected in blood. In a preliminary experiment, an average of 2 mosquitoes per mouse completed the meal (Supp Fig 1A) while an average of 4-6 engorged mosquitoes per mouse were collected for testing vaccine efficacy (Fig 1D). Keeping in mind that the number of engorged mosquitoes is only a fraction of the probing mosquitoes, and lethal infection was observed also in absence of complete meals, this suggests the lack of pathogenicity in this case was not dependent on an inadequate mosquito feeding attempt. In addition, PRVABC59 has been previously associated with loss of pathogenicity due to mutations acquired after a low number of passages in cell culture (35). This being said, in this study we confirmed ZIKV PRVABC59 lethality in AG129 mice in a challenge evaluation that compared side-by-side mosquito- and needle-mediated inoculation. In this specific instance, the higher susceptibility of the AG129 mouse strain is likely caused by the double knock-out genotype, however differences in host susceptibility also points towards the importance of the interactions between the host genetic background and the viral strain. Despite being associated with the most concerning human outbreaks, paradoxically ZIKV strains belonging to the Asian lineage show reduced *in vitro* replication and *in vivo* pathogenicity when compared to the African lineages (36, 37). Also, ZIKV strains belonging to the African lineage are reported to be more efficient in transmission than strains from Asian lineage (38). Taking into consideration all these variabilities, we tested the efficacy of our vaccine candidate using the most appropriate available vector-host transmission system, combining a commonly used mouse strain, a highly pathogenic ZIKV strain, such as the African ZIKV MP1751, and a highly susceptible lab-adapted *Ae. aegypti* mosquito strain having the globally invasive genotypic background.

While ZIKV infection can be sustained by sylvatic cycle, with non-human primates as a mammalian reservoir for the virus, densely human populated environments, combined with the remarkable anthropophilic behaviour of *Ae. aegypti*, well support ZIKV transmission in the urban cycle. In this study, allowing naïve mosquitoes to feed on viremic mice, we also assessed vaccine capacity to interfere with the transmission cycle, inhibiting the passage of the virus from vaccinated individuals to naïve mosquito vectors. A complete viral transmission with high titres was observed in mosquitoes fed on control non-vaccinated mice, while only one individual fed on a vaccinated mice showed a mild ZIKV infection in the carcass. When the potential of mosquitoes to transmit the virus was observed by analysing the titre of live viral particles in the SG, the vaccine was also able to efficiently interfere with mosquito capacity to spread the virus.

In conclusion, our findings confirm the capacity of the vaccine candidate VLP-cvD to protect mice against ZIKV infection also when the infection occurs through the bite of a mosquito vector. The vaccine is also capable of interfering with ZIKV spread by blocking the transmission of the virus from an infected mammalian host to uninfected mosquitoes. This study underlines the importance of considering the complexity of the vector-host-virus interaction when developing and evaluating disease control interventions, and further supports the potential of VLP-cvD as a vaccine candidate, making it worthy of further development for clinical end-use.

## Methods

### Animal ethics

all animal research described in this study was approved by the University of Glasgow Animal Welfare and Ethical Review Board and was carried out under United Kingdom Home Office Licenses, P9722FD8E, in accordance with the approved guidelines and under the UK Home Office Animals (Scientific Procedures) Act 1986 (ASPA).

### Cell lines and virus strain

Expi293F embryonic human kidney cells were maintained and transfected in Expi293(tm) Expression Medium (Thermo Fisher Scientific) as per the manufacturers’ protocol. Vero-furin cells (kindly supplied by Dr Theodore C. Pierson (39) were grown in Dulbecco’s modified Eagle’s medium (DMEM) (Life Technologies) containing 7% fetal bovine serum (FBS) (Life Technologies), 10 μg/ml of blasticidin (InvivoGen), and penicillin-streptomycin (Gibco). The virus was serially passaged in *Ae. albopictus* C6/36 cells: the infected supernatant was harvested, concentrated using Amicon Ultra-15 filters (Millipore, IRL), and titrated via fluorescent focus assay (FFA), as described below. ZIKV PRVABC59 (kindly supplied by BEI Resources; accession number KX087101) and ZIKV MP1751 (005V-02871; kindly supplied by Public Health England; accession number KY288905.1) were used for micro-neutralization, mosquito injections and animal challenges.

### VLP-cvD production and purification

VLP-cvD were produced in Expi293F cells at 28°C following transfection with a plasmid expressing ZIKV prM-E-A264C and purified as described in De Lorenzo et al (2020)(7) by performing, in sequence, 20% sucrose cushion, discontinuous 10-30% sodium potassium tartrate density gradient and size-exclusion chromatography with elution in PBS buffer. Eluted fractions were concentrated by ultrafiltration through Amicon® Ultra 15 (100 kDa, Merk Millipore).

### Mouse immunisation and ZIKV challenge

A129 mice or AG129KO mice (129Sv/Ev background; Marshall BioResources) at 4 weeks of age were immunised subcutaneously with either VLP-cvD or PBS, both adjuvanted with 1% AddaVax (InvivoGen). Needle challenge was performed by subcutaneous injection of PRVABC59 ZIKV (10^4^ pfu) or MP1751 ZIKV (100 pfu) in 100 µl 2%FBS-DMEM. Mosquito challenge was performed as described below. Pre- and post-challenge blood samples were collected for antibody titration, micro-neutralisation assay and serum viral load. After challenge, animal weight change and symptoms were daily monitored and scored for 14 days. Scoring system adapted was: 0 for no signs of distress or disease, 1 for one sign of distress, 2 for two signs of distress or mild disease, 3 for more than two signs of severe disease or loss of 15% percent of the body weight. Score of 3 was considered the human endpoint and mice were culled. Any individual mouse reaching clinical score of 3 or losing more than the 15% of the initial weight was euthanised. Surviving mice were euthanised at 14 days post challenge.

### ELISA for antibody titration

recombinant biotinylated sE-cvD protein was expressed at 28°C using an ExpiFectamine 293 transfection kit (Thermo Fisher Scientific). Cell supernatant was harvested and dialyzed. Biotinylated proteins were captured in ELISA plates precoated with 5 μg/ml of avidin (Sigma) in Na2CO3-NaHCO3 buffer (pH 9.6) and subsequently blocked with PBS containing 0.05% Tween 20 (PBST) and 1% bovine serum albumin (BSA; Sigma). Serial dilutions of mouse sera were tested for binding to the biotinylated proteins, and the bound antibodies were detected using horseradish peroxidase (HRP)-conjugated anti-mouse IgG A4416 (Sigma) and 3,3’,5,5’-tetramethylbenzidine (TMB) substrate (Life Technologies)

### Micro-neutralization assay

this assay was performed as described by Lopez-Camacho et al(11). Briefly,VERO-furin cells were seeded the day before the experiment at a density of 7 × 10^3^/well in 96-well plates. 3-fold serially diluted mice sera were first incubated at 37°C for 1 h with 100 PFU/well ZIKV. The serum/virus mix was then used to infect cells. After 1 h of incubation at 37°C, 100 μl of medium was added to each well. At day 3 post-infection, cells were lysed in lysis buffer (20 mM Tris-HCl [pH 7.4], 20 mM iodoacetamide, 150 mM NaCl, 1 mM EDTA, 0.5% Triton X-100, and cOmplete protease inhibitors), and the viral E protein quantitated by sandwich ELISA (see below). The amount of E protein detected correlates with the level of virus infectivity, which was presented as percentage of ZIKV infectivity relative to the control (i.e., virus not preincubated with immune sera). The MN50 titer was defined as the serum dilution that neutralized ZIKV infection by 50%.

### Sandwich ELISA for quantification of ZIKV infectivity

ELISA plates were coated with 3 μg/ml of purified pan-flavivirus monoclonal antibody (MAb) D1-4G2-4-15 (ATCC HB112TM) in PBS, incubated overnight at room temperature (RT), and blocked for 2 h at RT with PBST and 2% skimmed milk powder. After washing with PBST, ZIKV-infected cell lysates were added and incubated for 1 h at RT. Wells were washed with PBST, incubated with anti-ZIKV E rabbit polyclonal R34 IgG (9) at 6 μg/ml in PBST for 1 h at RT, then washed again. Antibodies bound to ZIKV envelope protein were detected using HRP-conjugated anti-rabbit IgG 7090 (Abcam) and TMB substrate (Life Technologies). The MN50 titre was defined as the serum dilution that neutralised >50% of ZIKV and was determine using Graphpad Prism 9, and nonlinear regression (curve fit) performed for the data points using Log (inhibitor) versus response (variable slope).

### Mosquito rearing

the *Ae. aegypti* wild-type line used was colonized from Selangor State, Malaysia in the 1960s. Colonies were maintained in standard rearing conditions, at 27°C and 70% relative humidity with a 12-hour light/dark cycle. Larvae were fed on tropical fish pellets (Tetramin, Tetra, Melle, Germany) and adults maintained with 5% sucrose solution *ad libitum*. Blood meals to maintain the colony were provided using an artificial blood-feeding system (Hemotek, UK) using human blood (Scottish National Blood Transfusion Service, UK). Eggs were collected on a wet filter-paper (Grade 1 filter paper, Whatman plc, GE healthcare, UK), desiccated for 5 days and later hatched in deionized water containing 1g/L bovine liver powder (MP Biomedicals, Santa Ana, California, USA).

### Mosquito infection

5-day old female mosquitoes were infected by intra-thoracically injection of 10^6^ FFU/ml of ZIKV using a Nanoject II (Drummond Scientific, USA) hand-held microinjector. Injected mosquitoes were immediately transferred into a climatic chamber at 27°C, with 70% relative humidity and a 12-hour light/dark cycle for recovery. ZIKV-injected females were maintained on sugar solution for 7 days prior mice infection.

### Mosquito feeding on mice

mice were transferred from the animal house to the insectary where they were anesthetized using a Ketamine-Medetomidine cocktail (Domitor®) based on their body weight (Ketamine 50 mg/kg -Medetomidine 0.5 mg/kg). Once anaesthesia was effective, each mouse was transferred for 20 minutes on the organdie cover of a mosquitoes-containing cardboard cup, with the eyes protected while the mosquitoes fed through the net. At the end of the feeding, the anaesthetic effects were reversed with Atipamezole (Antisedant)(1 mg/kg) and mice condition monitored for the rest of the day. After feeding, the number of fully engorged mosquitoes was recorded for every mouse. Cardboard cups with fed mosquitoes were placed back in the climatic chambers.

### RT-qPCR for mouse viremia

viral RNA was extracted from post-challenge sera by QIAamp® viral RNA mini kit (QIAGEN). Viral load was measured by RT-qPCR using One-Step SYBR® Primescript(tm) RT-PCR kit II (Takara). CT values from serum samples were used to calculate serum viral load according to regression equation built by a set of standard viral RNA extracted from dilutions of known titre virus preparation. Primers pair for PRVABC59 ZIKV gene was Forward:5’-TTGGTCATGATACTGCTGATTGC -3’ and Reverse: 5’-CCTTCCACAAAGTCCCTATTGC -3’, while for MP1751 ZIKV gene, we used Forward:5’-ACTTCCGGTGCGTTACATGA-3’ and Reverse:5’-GGGCTTCATCCATGATGTAG-3’.

### RT-qPCR for mosquitoes

RNA was extracted using TRI Reagent (Sigma-Aldrich, Missouri, USA). cDNA was synthesized using 1μg of total RNA and the All-In-One cDNA Synthesis SuperMix (Biotool, Houston, Texas, USA). qRT-PCRs were performed on a 1 to 20 dilution of the cDNAs. Virus levels were normalized to the RpS17 house-keeping gene (RpS17-F 5’-CACTCCCAGGTCCGTGGTAT-3’; RpS17-R: 5’-GGACACTTCCGGCACGTAGT-3’)

### Viral quantification in mosquito salivary glands

mosquito salivary glands were dissected and sampled in Dulbecco’s Modified Eagle Medium (DMEM) supplemented with 2% fetal bovine serum (FBS). After homogenizing, 10-fold serial dilutions of each solution were transferred onto a monolayer of Vero-furin cells for viral quantification with FFA. Primary antibody for Zika was a Mouse Monoclonal antibody DIII1B (7); secondary antibody was the Goat anti-mouse Alexa Fluor 488, A-11001 (Thermo Scientific, Waltham, Massachusetts, USA). Celigo Imaging Cytometer (Nexcelom Bioscience, Lawrence, Massachusetts) was used for imaging plates. Fluorescent foci were quantified by eye (from dilutions with less than 100 foci) and virus titers calculated and expressed as FFU/ml.

### Statistical analysis

all graphs and statistical analyses were produced using GraphPad Prism 9 (GraphPad Software Inc., San Diego, CA, USA). A Shapiro-Wilk test was used for assessing normality distribution of data, and parametric and nonparametric tests were selected accordingly for antibody and virus titre. Multiple comparisons were calculated using Bonferroni method for p values adjustment. Statistical analysis of survival was performed using a Log-rank (Mantel-Cox) test with 95% confidence interval. Statistical analysis of viremia was performed using a 2-sided analysis of variance (ANOVA), with Tukey’s pairwise comparison at 95% confidence interval. 0.0332 (*), 0.0021 (**), 0.0002 (***), <0.0001 (****).

## Acknowledgments

we acknowledge the assistance of Nicola Munro, Catrina Boyd, Stuart Lannigan and Scott McCall at Biological Services, University of Glasgow. We are grateful to Dr Theodore C. Pierson for kindly suppling the VERO-furin cell line. We acknowledge the provision of ZIKV strains PRVABC59 and MP1751 (005V-02871) from BEI Resources and Public Health England/EVAg, respectively. This research was funded by the Department of Health and Social Care using UK Aid funding and is managed by the NIHR, and by the UK Medical Research Council, MC_UU12014/2 for AHP and MC_UU12014/8 for AK. The views expressed in this publication are those of the authors and not necessarily those of the Department of Health and Social Care. This project was also partially funded through the Wellcome Trust award 202888/Z/16/Z (SPS). The funders had no role in study design, data collection and analysis, the decision to publish, or preparation of the manuscript.

## Author contributions

Authors’ contributions are as follows. Conceptualization, MVM, GDL; Methodology MVM, RT, THA, DDG, SMM, CS, PN, GDL; Validation, MVM, RT, THA, DDG, GDL; Formal analysis, MVM, RT, THA, DDG, GDL; Investigation, MVM, RT, THA, DDG, CS, PN, GDL; Resources, AK, SPS, AHP; Data curation, MVM, RT, THA, DDG, GDL; Writing – Original Draft, MVM, GDL; Writing – Review & Editing, MVM, RT, THA, DDG, SMM, CS, PN, AK, SPS, AHP, GDL; Visualization, MVM, GDL; Supervision, MVM, GDL; Project Administration, MVM, AHP, GDL; Funding Acquisition, SPS, AHP, GDL.

---

**Supplementary Figure 1:**
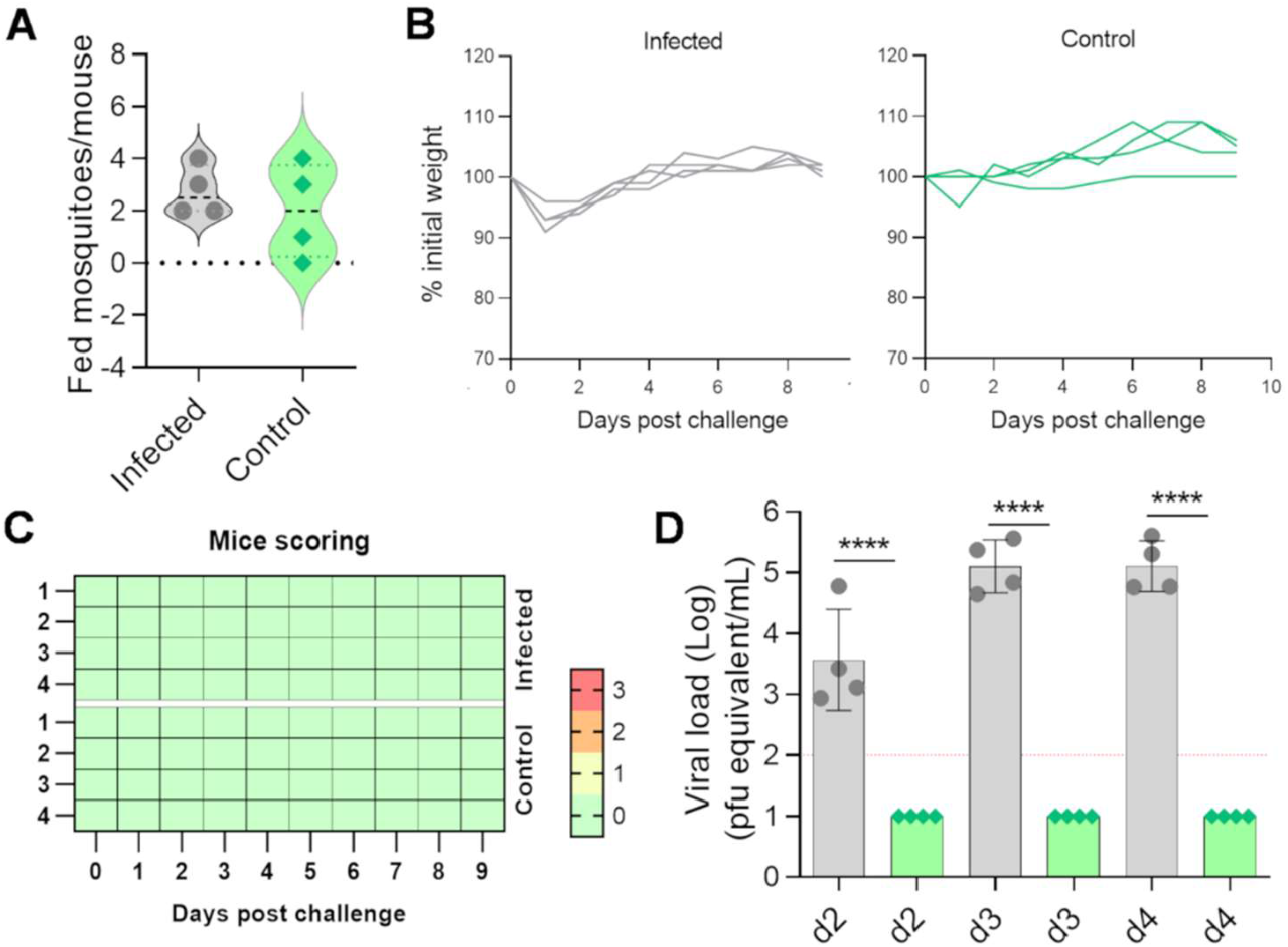
**A)** Number of fed mosquitoes per mouse at the end of the feeding procedure. Grey indicates PRVABC59 infected mosquitoes, green indicates naïve mosquitoes. **B)** Animal body weight variations, calculated as percentage of the initial weight. Clinical scoring for signs of infection. **D)** Viral titre in challenged animals at days 2, 3 and 4 post infection. Columns show geometric means from all mice with geometric SD. Dotted line indicates limit of detection.

**Supplementary Figure 2:**
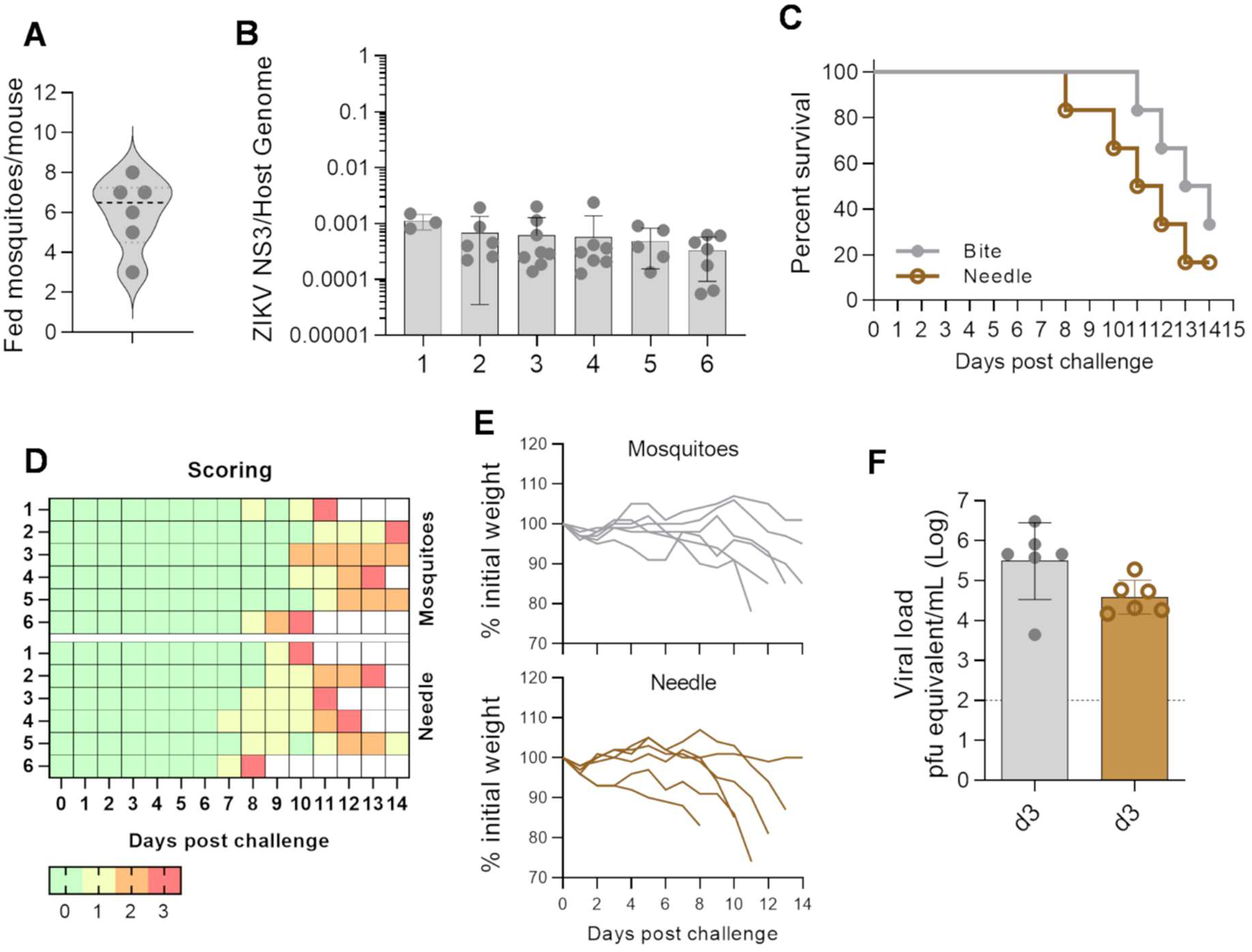
**A)** Number of fed mosquitoes per mouse at the end of the feeding procedure. **B** Relative quantification of ZIKV genome copies normalized on the host genome from whole mosquito bodies. **C)** Percentage of survival mice during the 14-days challenge. Grey indicates mosquito bite-mediated infection, brown indicates needle-mediated infection. Clinical scoring for signs of infection. **E)** Mice body weight variations, calculated as percentage of the initial weight. **F)** Viral titre in challenged animals at day 3 post-challenge. Columns show geometric means from all mice with geometric SD. Dotted line indicates limit of detection.

## Bibliography

1. Munoz LS, Barreras P, Pardo CA. Zika Virus-Associated Neurological Disease in the Adult: Guillain-Barre Syndrome, Encephalitis, and Myelitis. Semin Reprod Med. 2016;34(5):273–9.

2. Oehler E, Watrin L, Larre P, Leparc-Goffart I, Lastere S, Valour F, et al. Zika virus infection complicated by Guillain-Barre syndrome--case report, French Polynesia, December 2013. Euro Surveill. 2014;19(9).

3. Mlakar J, Korva M, Tul N, Popovic M, Poljsak-Prijatelj M, Mraz J, et al. Zika Virus Associated with Microcephaly. N Engl J Med. 2016;374(10):951–8.

4. Vhp L, Aragao MM, Pinho RS, Hazin AN, Paciorkowski AR, Penalva de Oliveira AC, et al. Congenital Zika Virus Infection: a Review with Emphasis on the Spectrum of Brain Abnormalities. Curr Neurol Neurosci Rep. 2020;20(11):49.

5. D’Ortenzio E, Matheron S, Yazdanpanah Y, de Lamballerie X, Hubert B, Piorkowski G, et al. Evidence of Sexual Transmission of Zika Virus. N Engl J Med. 2016;374(22):2195–8.

6. Ades AE, Thorne C, Soriano-Arandes A, Peckham CS, Brown DW, Lang D, et al. Researching Zika in pregnancy: lessons for global preparedness. Lancet Infect Dis. 2020;20(4):e61–e8.

7. De Lorenzo G, Tandavanitj R, Doig J, Setthapramote C, Poggianella M, Sanchez-Velazquez R, et al. Zika Virus-Like Particles Bearing a Covalent Dimer of Envelope Protein Protect Mice from Lethal Challenge. J Virol. 2020;95(1).

8. Richner JM, Himansu S, Dowd KA, Butler SL, Salazar V, Fox JM, et al. Modified mRNA Vaccines Protect against Zika Virus Infection. Cell. 2017;169(1):176.

9. Cimica V, Galarza JM, Rashid S, Stedman TT. Current development of Zika virus vaccines with special emphasis on virus-like particle technology. Expert Rev Vaccines. 2021;20(11):1483–98.

10. Gaudinski MR, Houser KV, Morabito KM, Hu Z, Yamshchikov G, Rothwell RS, et al. Safety, tolerability, and immunogenicity of two Zika virus DNA vaccine candidates in healthy adults: randomised, open-label, phase 1 clinical trials. Lancet. 2018;391(10120):552–62.

11. Lopez-Camacho C, Abbink P, Larocca RA, Dejnirattisai W, Boyd M, Badamchi-Zadeh A, et al. Rational Zika vaccine design via the modulation of antigen membrane anchors in chimpanzee adenoviral vectors. Nat Commun. 2018;9(1):2441.

12. Morrison TE, Diamond MS. Animal Models of Zika Virus Infection, Pathogenesis, and Immunity. J Virol. 2017;91(8).

13. Routhu NK, Lehoux SD, Rouse EA, Bidokhti MRM, Giron LB, Anzurez A, et al. Glycosylation of Zika Virus is Important in Host-Virus Interaction and Pathogenic Potential. Int J Mol Sci. 2019;20(20).

14. Talavera-Aguilar LG, Murrieta RA, Kiem S, Cetina-Trejo RC, Baak-Baak CM, Ebel GD, et al. Infection, dissemination, and transmission efficiencies of Zika virus in Aedes aegypti after serial passage in mosquito or mammalian cell lines or alternating passage in both cell types. Parasit Vectors. 2021;14(1):261.

15. Styer LM, Kent KA, Albright RG, Bennett CJ, Kramer LD, Bernard KA. Mosquitoes inoculate high doses of West Nile virus as they probe and feed on live hosts. PLoS Pathog. 2007;3(9):1262–70.

16. Schneider BS, Higgs S. The enhancement of arbovirus transmission and disease by mosquito saliva is associated with modulation of the host immune response. Trans R Soc Trop Med Hyg. 2008;102(5):400–8.

17. Styer LM, Lim PY, Louie KL, Albright RG, Kramer LD, Bernard KA. Mosquito saliva causes enhancement of West Nile virus infection in mice. J Virol. 2011;85(4):1517–27.

18. Moser LA, Lim PY, Styer LM, Kramer LD, Bernard KA. Parameters of Mosquito-Enhanced West Nile Virus Infection. J Virol. 2016;90(1):292–9.

19. McCracken MK, Gromowski GD, Garver LS, Goupil BA, Walker KD, Friberg H, et al. Route of inoculation and mosquito vector exposure modulate dengue virus replication kinetics and immune responses in rhesus macaques. PLoS Negl Trop Dis. 2020;14(4):e0008191.

20. Le Coupanec A, Babin D, Fiette L, Jouvion G, Ave P, Misse D, et al. Aedes mosquito saliva modulates Rift Valley fever virus pathogenicity. PLoS Negl Trop Dis. 2013;7(6):e2237.

21. Jin L, Guo X, Shen C, Hao X, Sun P, Li P, et al. Salivary factor LTRIN from Aedes aegypti facilitates the transmission of Zika virus by interfering with the lymphotoxin-beta receptor. Nat Immunol. 2018;19(4):342–53.

22. Schneider BS, Soong L, Coffey LL, Stevenson HL, McGee CE, Higgs S. Aedes aegypti saliva alters leukocyte recruitment and cytokine signaling by antigen-presenting cells during West Nile virus infection. PLoS One. 2010;5(7):e11704.

23. Wanasen N, Nussenzveig RH, Champagne DE, Soong L, Higgs S. Differential modulation of murine host immune response by salivary gland extracts from the mosquitoes Aedes aegypti and Culex quinquefasciatus. Med Vet Entomol. 2004;18(2):191–9.

24. Boppana VD, Thangamani S, Adler AJ, Wikel SK. SAAG-4 is a novel mosquito salivary protein that programmes host CD4 T cells to express IL-4. Parasite Immunol. 2009;31(6):287–95.

25. Pingen M, Bryden SR, Pondeville E, Schnettler E, Kohl A, Merits A, et al. Host Inflammatory Response to Mosquito Bites Enhances the Severity of Arbovirus Infection. Immunity. 2016;44(6):1455–69.

26. McCracken MK, Christofferson RC, Grasperge BJ, Calvo E, Chisenhall DM, Mores CN. Aedes aegypti salivary protein “aegyptin” co-inoculation modulates dengue virus infection in the vertebrate host. Virology. 2014;468-470:133-9.

27. Pingen M, Schmid MA, Harris E, McKimmie CS. Mosquito Biting Modulates Skin Response to Virus Infection. Trends Parasitol. 2017;33(8):645–57.

28. Schmid MA, Glasner DR, Shah S, Michlmayr D, Kramer LD, Harris E. Mosquito Saliva Increases Endothelial Permeability in the Skin, Immune Cell Migration, and Dengue Pathogenesis during Antibody-Dependent Enhancement. PLoS Pathog. 2016;12(6):e1005676.

29. Thangamani S, Higgs S, Ziegler S, Vanlandingham D, Tesh R, Wikel S. Host immune response to mosquito-transmitted chikungunya virus differs from that elicited by needle inoculated virus. PLoS One. 2010;5(8):e12137.

30. Surasombatpattana P, Ekchariyawat P, Hamel R, Patramool S, Thongrungkiat S, Denizot M, et al. Aedes aegypti saliva contains a prominent 34-kDa protein that strongly enhances dengue virus replication in human keratinocytes. J Invest Dermatol. 2014;134(1):281–4.

31. Dudley DM, Newman CM, Lalli J, Stewart LM, Koenig MR, Weiler AM, et al. Infection via mosquito bite alters Zika virus tissue tropism and replication kinetics in rhesus macaques. Nat Commun. 2017;8(1):2096.

32. Peters NC, Kimblin N, Secundino N, Kamhawi S, Lawyer P, Sacks DL. Vector transmission of leishmania abrogates vaccine-induced protective immunity. PLoS Pathog. 2009;5(6):e1000484.

33. Lopez-Camacho C, De Lorenzo G, Slon-Campos JL, Dowall S, Abbink P, Larocca RA, et al. Immunogenicity and Efficacy of Zika Virus Envelope Domain III in DNA, Protein, and ChAdOx1 Adenoviral-Vectored Vaccines. Vaccines (Basel). 2020;8(2).

34. Dowall SD, Graham VA, Rayner E, Hunter L, Atkinson B, Pearson G, et al. Lineage-dependent differences in the disease progression of Zika virus infection in type-I interferon receptor knockout (A129) mice. PLoS Negl Trop Dis. 2017;11(7):e0005704.

35. Duggal NK, McDonald EM, Weger-Lucarelli J, Hawks SA, Ritter JM, Romo H, et al. Mutations present in a low-passage Zika virus isolate result in attenuated pathogenesis in mice. Virology. 2019;530:19–26.

36. Aubry F, Jacobs S, Darmuzey M, Lequime S, Delang L, Fontaine A, et al. Recent African strains of Zika virus display higher transmissibility and fetal pathogenicity than Asian strains. Nat Commun. 2021;12(1):916.

37. Simonin Y, van Riel D, Van de Perre P, Rockx B, Salinas S. Differential virulence between Asian and African lineages of Zika virus. PLoS Negl Trop Dis. 2017;11(9):e0005821.

38. Aubry F, Dabo S, Manet C, Filipovic I, Rose NH, Miot EF, et al. Enhanced Zika virus susceptibility of globally invasive Aedes aegypti populations. Science. 2020;370(6519):991–6.

39. Mukherjee S, Sirohi D, Dowd KA, Chen Z, Diamond MS, Kuhn RJ, et al. Enhancing dengue virus maturation using a stable furin over-expressing cell line. Virology. 2016;497:33-40.

